# Inferential Pitfalls in Decoding Neural Representations

**DOI:** 10.1101/141283

**Authors:** Vencislav Popov, Markus Ostarek, Caitlin Tenison

**Author notes:** VP and MO contributed equally and should be considered co-first authors.

## Abstract

A key challenge for cognitive neuroscience is to decipher the representational schemes of the brain. A recent class of decoding algorithms for fMRI data, stimulus-feature-based encoding models, is becoming increasingly popular for inferring the dimensions of neural representational spaces from stimulus-feature spaces. We argue that such inferences are not always valid, because decoding can occur even if the neural representational space and the stimulus-feature space use different representational schemes. This can happen when there is a systematic mapping between them, as shown by two simulations. In one simulation, we successfully decoded the binary representation of numbers from their decimal features. Since binary and decimal number systems use different representations, we cannot conclude that the binary representation encodes decimal features. In the second simulation, we successfully decoded the HSV color representation from the RGB representation of colors, even though these color spaces have different geometries and their dimensions have different interpretations. Detailed analysis of the predicted colors showed systematic deviations from the ground truth despite the high decoding accuracy, indicating that decoding accuracy on its own is not sufficient for making representational inferences. The same argument applies to the decoding of neural patterns from stimulus-feature spaces and we urge caution in inferring the nature of the neural code from such methods. We discuss ways to overcome these inferential limitations.

## I. Introduction

A key challenge for cognitive neuroscience is to decipher the representational schemes of the brain, that is, to understand the neural code that underlies the encoding and representation of sensory, motor, spatial, emotional, semantic and other types of information. To address these issues, researchers often employ neuroimaging techniques like functional magnetic resonance imaging (fMRI), which measures the blood oxygenation level-dependent (BOLD) activation in the brain that is elicited when participants engage with different stimuli. A common assumption has been that the underlying neural representation of each stimulus has measurable but complex effects on the BOLD activation patterns. In order to understand what those patterns of activity can tell us about how the brain processes and represents information, researchers have used various analytical tools such as univariate subtraction methods, multivariate pattern (MVP) classification, representational similarity analysis (RSA) and, recently, explicit stimulus-feature-based encoding and decoding models (for reviews, see Davis & Poldrack, 2013, Haxby, Connolly, & Guntupalli, 2014, or Naselaris, Kay, Nishimoto, & Gallant, 2011). Despite their differences, all of these methods have the same goal – to quantify how changes in task conditions and the properties of the stimuli relate to changes in BOLD activation and vice versa. One way in which these methods differ is in how they achieve that mapping and in what inferences they allow us to draw.

In this article, we review some of the known inferential limitations of existing fMRI analysis methods and we highlight a previously unrecognized issue in interpreting results from stimulus-feature-based encoding and decoding models. The latter are steadily becoming the de facto gold standard for investigating neural representational spaces (Haxby et al. 2014, Naselaris & Kay, 2015).

## II. Univariate vs. multivariate analysis

Two of the main questions that any fMRI analysis technique attempts to answer are 1) where information is represented/processed and 2) what is the nature of those representations (Davis & Poldrack, 2013). The localization and the representational questions are related, but distinct, and one commonality between all methods we review below is that understanding *how* information is represented turns out to be more difficult that understanding *where* it is represented or processed.

This difference is most easily exemplified in univariate subtraction analyses. Before the advent of the more advanced techniques we review below, the main fMRI analysis tool was based on comparing how activity in a single voxel or averaged activity in a contiguous area of voxels differs between task conditions or stimuli (see Figure 1). Researchers have used these univariate subtraction methods successfully to understand the relative engagement of certain brain areas in specific tasks. For example, the observation that subsequently remembered stimuli cause greater hippocampal activation during their *encoding* compared to subsequently forgotten stimuli, has corroborated conclusions from lesion studies that the medial temporal lobe is somehow involved in memory formation (Wagner et al., 1998). Unfortunately, the coarse nature of this method precludes fine-grained inferences about the underlying representational content and computations that give rise to the observed BOLD signal. Univariate methods assume that representational or processing differences between stimuli can be observed in individual voxels, and they ignore any relationships between different voxels (Davis & Poldrack, 2013). By ignoring the possibility that information might be represented in a distributed manner across voxels, the assumptions underlying univariate subtraction methods limit their use in understanding neural representations. In addition, these methods cannot tell us whether changes in activation are due to representational preferences, processing differences, or attentional variation among conditions (Coutanche, 2013).

**Figure 1.**
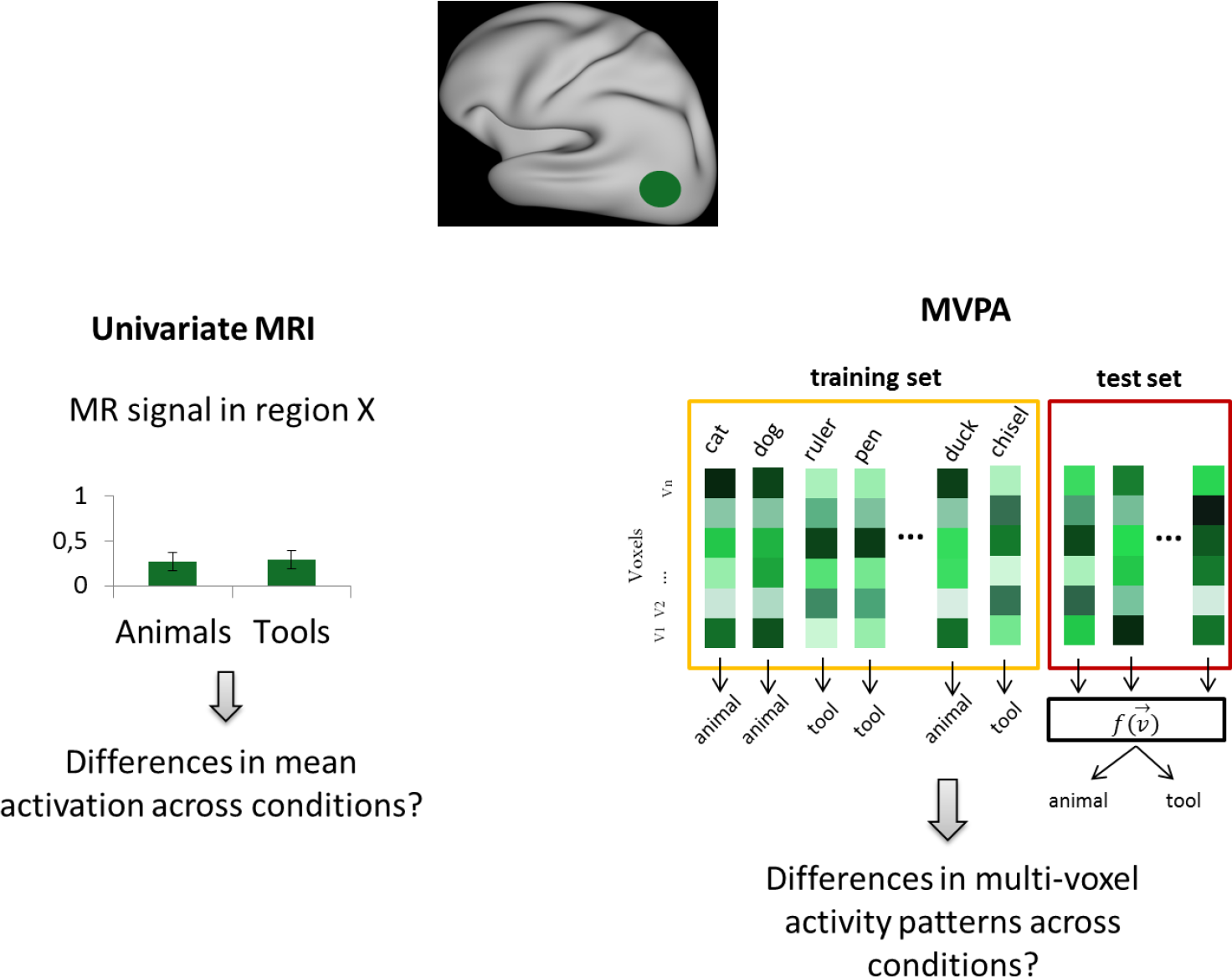
Univariate vs. Multivariate fMRI. In univariate fMRI (left), the average activation level in a brain region is compared across experimental conditions (e.g., animal vs. tool words). In multivoxel pattern analysis (right), machine learning techniques are used to train a classifier to distinguish categories based on fine-grained activation patterns (yellow box) and its classification performance is evaluated on a separate test set (red box).

In contrast, multivoxel pattern analysis (MVPA) techniques have attempted to overcome this limitation by looking at how various categories of stimuli or task conditions lead to combinatorial differences (i.e. MVP classification) or similarities (i.e. RSA, see Figure 2) in distributed patterns of activity over multiple voxels. These methods have become popular because they allow researchers to study neural representational spaces with increasing sensitivity and resolution. For example, a seminal study by Haxby et al. (2001) found that visual object categories can be classified based on the pattern of activation that their exemplars elicited in the ventral temporal cortex. The classification was successful despite the lack of overall activation differences in that region. Similar methods have been used to show that concepts have language-invariant representations in the anterior temporal lobe (Correia et al., 2014), that very similar visual scenes can be discriminated in the hippocampus (Bonnici et al., 2012) and that during their retrieval from memory, the shape, color and identity of visual objects can be differentially decoded across several cortical areas (Coutanche & Thompson-Schill, 2015).

**Figure 2.**
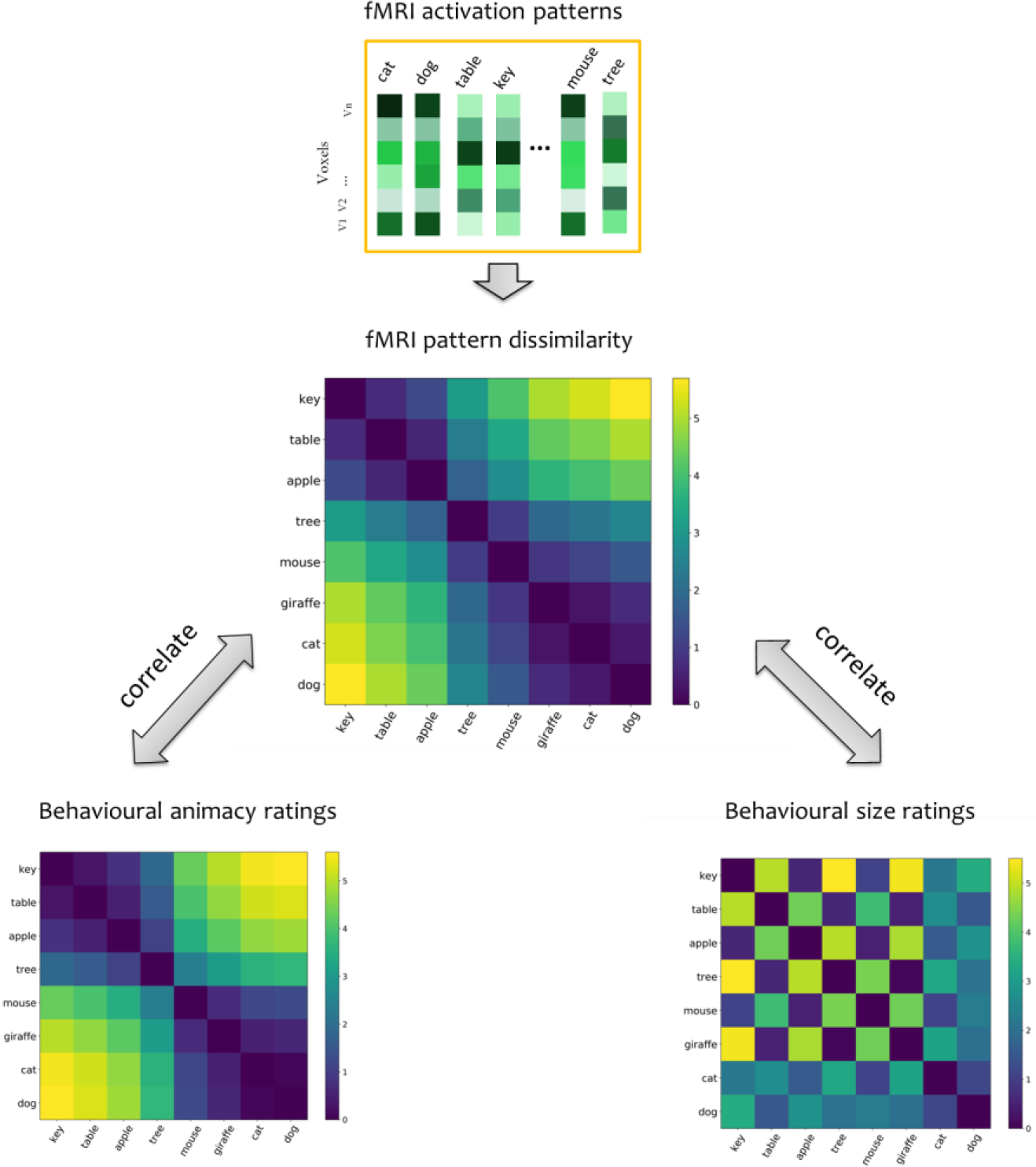
Representational Similarity Analysis (RSA). Multivoxel activation patterns of all items or conditions (e.g., words) are estimated (top panel) and compared to each other (typically by calculating 1 - correlation) to produce a representational dissimilarity matrix (RDM; middle panel) that abstracts away from the precise patterns related to each item and instead quantifies their dissimilarity. RDMs derived from neural activation patterns can be correlated with model RDMs that reflect hypotheses about the organization of informational content in a given brain region. In this hypothetical example, the RDM based on activity patterns strongly correlates with the RDM derived from behavioral ratings of animacy, but not with the RDM derived from behavioral ratings of size.

Despite early enthusiasm that MVPA methods could be used to understand the structure of the neural code and the nature of the underlying representations (Norman, Polyn, Detre, & Haxby, 2006), conventional MVP classification and RSA techniques share one of the same fundamental inferential limitations of univariate methods. Successful classification or careful inspection of confusions/similarity matrices can indicate that some relevant information about the stimulus class is present in the population of analyzed voxels, thus answering the *where question*, but it cannot identify exactly what that information is, or how it is represented and organized (Naselaris & Kay, 2015; Poldrack, 2011; Tong & Pratte, 2012). Because neural data is correlational, many different properties of the stimuli might lead to successful classification of the stimulus category, the task condition, or the brain state in question. For example, successfully categorizing whether a word represents an animate or an inanimate object does not necessarily mean that the region of interest encodes that category distinction. There are many differences between animate and inanimate objects, such as differences in their sensory and functional features (Farah & McClelland, 1991) that could be responsible for the successful classification.

In a similar argument, Ritchie, Kaplan, and Klein (2017) recently pointed out that the main reason behind this limitation is that linear classifiers are "informationally greedy": they are likely to pick up any information that distinguishes the to-be-classified categories and it is therefore often impossible to know what information they use. For instance, it remains unclear whether the orientation of a grating can be decoded from visual cortex because imperfect sampling of orientation-selective columns leads to hyperacuity (Kamitani & Tong, 2005), because of radial biases in the retinotopic map (Freeman, 2011), or because of edge-related activity (Carlson, 2014). Ritchie et al. (2017) further suggest that relating classifier performance to behavior only partly remedies the problem as “a brain region might carry information which is reliably correlated with the information that is actually used, but which is not itself used in behavior".

Another limitation of conventional MVP classifiers is that they cannot generalize and predict behavioral responses to novel *types* of stimuli or task conditions. To understand why, we can conceptualize classifiers in terms of types and tokens. An MVP classifier is usually trained on stimuli that are tokens from several types. The stimuli tokens might be different category exemplars, and the classifier is trained to predict the type of category to which they belong. Alternatively, the tokens might be multiple presentations of the same word in different modalities or languages and the types are the unique words themselves. In the first case, the classifier can only be used to predict category membership of items that belong to one of the categories on which it was trained. For instance, if one trains a classifier to predict the color of objects and trains it on yellow and orange objects (Coutanche & Thompson-Schill, 2015), one will not be able to predict the color of novel objects that are green. In the second case, even though the classifier could be used to predict exemplars in novel languages or modalities, it is again restricted only to exemplars of the words on which it was trained in the first place. In general, while the tokens being tested might be novel, they will be potentially decoded only if they are exemplars of a type that has already been trained on.

This methodological limitation is important - just as understanding how the decimal system represents numbers allows people to understand and manipulate numbers they have never seen before, a complete understanding of any neural representational system should allow researchers to use the neural pattern associated with novel stimuli to predict their identity, even if those stimuli are not exemplars of the types on which a particular model was trained on.

## III. Encoding models

To overcome these limitations many researchers are turning to a novel analysis method that is known by a few different names – voxelwise modelling (Naselaris & Kay, 2015), stimulus-model based encoding and decoding (Haxby et al., 2014), voxel-based encoding and decoding models (Naselaris et al., 2011), and forward models (Brouwer & Heeger, 2009; Fernan-dino, Humphries, Conant, Seidenberg, & Binder, 2016). For simplicity, we will refer to this class of methods as *encoding models* for the rest of this article. Encoding models can decode the identity of novel *types* of stimuli from neural activity by predicting activity not for the stimuli themselves, but for a set of simpler features into which they can be decomposed (see Figure 3). In a seminal study, Mitchell et al. (2008) predicted the neural activity associated with individual novel words based only on the activation of other words. To achieve that, they decomposed each word into a vector of weights on 25 sensory-motor semantic features (verbs such as “eat”, “taste”, “run”, “fear”, etc.). The weights were estimated from co-occurrence statistics of the word with each verb feature in a large corpus. They trained a classifier to predict the neural activity associated with *each constituent feature* of a training set of words, which resulted in separate neural activation maps for each feature. Neural activity for novel test words was then predicted highly accurately as a linear combination of the semantic feature activation maps weighted by the association of the word with each feature. Based on these results, Mitchell et al. (2008) concluded that the neural representation of concrete nouns might be based on sensory-motor features.

**Figure 3.**
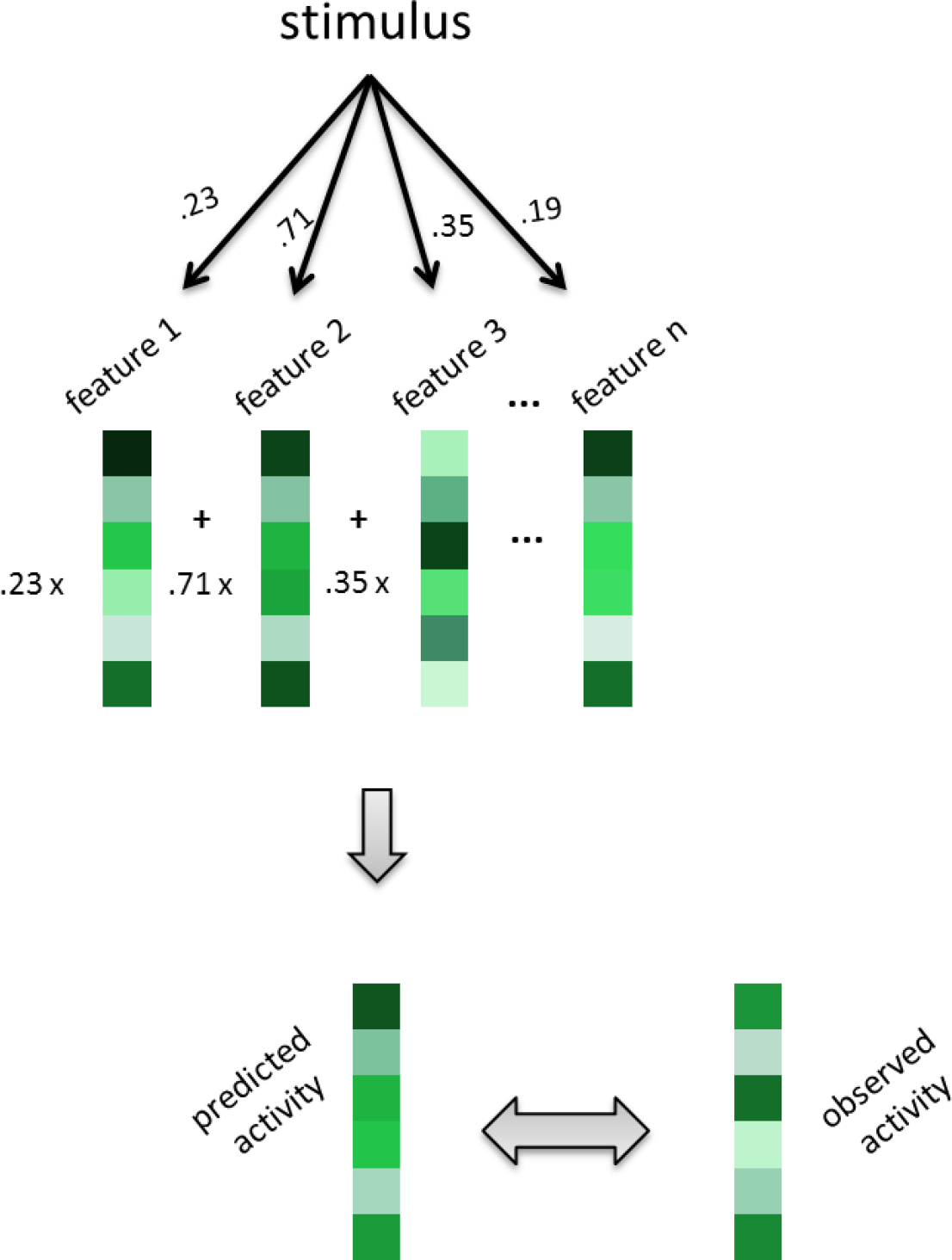
Encoding models. Stimuli are broken down into a set of features that are hypothesized to underlie stimulus representation. During training, a model learns to associate each feature's weights with voxel-wise activity, resulting in one feature activation maps per feature. The model then predicts voxel-wise activity to unseen test stimuli as the linear combination of these feature activation maps weighted by the test items' association with each feature. Predicted activity is compared to the actually observed activity per voxel to quantify prediction accuracy.

Similar approaches have been used to predict the neural response to novel natural images using Gabor filter features (Kay, Naselaris, Prenger, & Gallant, 2008), to novel colors based on color tuning curve features (Brouwer & Heeger, 2009), to novel music clips based on acoustic timbre features (Casey, Thompson, Kang, Raizada, & Wheatley, 2012), to natural sounds based on frequency, spectral and temporal modulations (Santoro et al., 2014), to novel faces based on a PCA decomposition of face features (Lee & Kuhl, 2016), to novel words based on subjective sensory-motor ratings (Fernandino et al., 2016). The motivating question behind many of these studies has been about the nature of the representations used by the brain in encoding the experimental stimuli, and the results have been used to argue that the neural representation is based on the constituent features of the stimuli used in the model.

In general, most encoding models use the following analysis procedure (see also Figure 3):

1. Specify a set of features that hypothetically underlie the representation of a stimulus set in the brain
2. Decompose a set of stimuli into vectors of weights for each feature
3. Select a region of interest (ROI) in the brain from which to analyze neural activation
4. Train a model to predict activity in each voxel for a training set of stimuli, using the weights of their features as predictors
5. Derive activation pattern maps (e.g. regression coefficients) associated with each feature
6. Predict neural activity in the ROI for novel stimuli, based on their feature weights and the activation pattern maps for each feature
7. Compare predicted neural activity for each novel stimulus with their observed neural activity and derive a measure of fit and accuracy

Thus, encoding models attempt to map a stimulus feature representational space, where each feature is a separate dimension, and each stimulus is a point in that space, to a neural activation space, where each voxel or each neuron is a separate dimension, and the activation pattern elicited by each stimulus is a point in that space (Kriegeskorte & Kievit, 2013).

Such representational models can be established either at the neuronal level, voxel level or both. Voxels are not the computational units of the brain, and they only indirectly reflect the averaged activity of hundreds of thousands of neurons (Henson, 2005). However, in most cases researchers do not have the ability to record large-scale neuronal activity directly in human participants. For that reason, some encoding models specify not only how individual neurons would respond to certain stimulus features, but also how the averaged activity of neurons within a voxel would translate to voxel-level BOLD activation. For example, Brouwer & Heeger (2009) studied how colors are represented in the visual cortex. In building their encoding model, they assumed that neurons in V4 have tuning curves for each of 6 distinct hues, and that the hue of a color is represented with a population code over these 6 types of neurons. They also assumed that the 6 types of neurons are not distributed uniformly in V4, such that each voxel contains different proportions of each type of neuron. Their encoding model estimated the proportion of each neuron type within each voxel by training the model on 7 out of 8 color types, and then used these proportions as weights to predict neural activity for the eight color type, which was never presented to the model during training. Thus, their encoding model specified on one level how individual neurons represent hue values, and then specified on a second level how the neuronal activity is combined to produce the observable voxel activations.

This two-level encoding model approach was also recently extended to the analysis of electroencephalography (EEG) data (Foster, Sutterer, Serences, Vogel, & Awh, 2016). In an EEG study on spatial working memory, Foster et al. (2016) showed that they can decode the spatial location of items held in working memory based on the topographic distribution of alpha-band oscillatory power. Analogously to Brouwer & Heeger (2009) approach to fMRI, Foster et al. (2016) assumed that each electrode reflects a weighted sum of the activation of several types of neurons, called spatial channels, where each channel had a receptive field for a different angular location. These angular channels were assumed to have a graded response, being most active for their preferred angular location, and progressively less active for other locations. The authors showed that the reconstructed activity of each channel from the alpha-band power closely tracks the spatial location currently maintained in working memory.

While this population coding approach is popular in studying perceptual representations (Brouwer & Heeger, 2009; Foster, Sutterer, Serences, Vogel, & Awh, 2016; Kay, Naselaris, Prenger, & Gallant, 2008), due to the relatively simple and better understood organization of the sensory cortices, it is rarely applied in studies on semantic representations. To date, there are no detailed proposals of how individual neurons and population of neurons represent semantic features and dimensions, nor what the tuning curves of those neurons might be. For this reason, researchers who study the nature of semantic representations with encoding models have so far specified more generic representational models at the voxel level and have made no claims about the nature of the underlying neuronal population code (e.g., Fernandino et al., 2017; Huth, Nishimoto, Vu, & Gallant, 2012; Mitchel et al., 2008). Undoubtedly, as our understanding of semantic representational spaces grows, bridging the neural-to-voxel representational gap would become increasingly important.

## IV. What can we infer about neural representations?

As our brief review indicated, encoding models have grown in popularity due to their many advantages over more classical classification approaches. While there is little doubt that these models are a useful and powerful analytic tool, their growing use requires us to carefully consider what we can and cannot conclude on the basis of successful decoding. What can a successful mapping between a stimulus feature space and a neural activation space tell us about the nature of the representation used by the brain? A common inference in some of these studies has been that if you can predict the identity of novel stimuli based on that mapping, then the neural representation is likely based on the feature set used by the model. Put formally, the inferential claim goes as follows:

- We can represent certain stimuli as a combination of lower-level features
- We can show that it is possible to predict the neural pattern caused by a novel stimulus in brain area A from an encoding model based on these features
- *Therefore, brain area A encodes those features and uses a representational scheme based on them.*

This claim has been made to different degrees both in theoretical and methodological papers reviewing the approach (e.g., Haxby et al., 2014; Naselaris & Kay, 2015; Naselaris et al., 2011; Norman et al., 2006; Ritchie, Kaplan, & Klein, 2017; Tong & Pratte, 2012), as well as in empirical studies that use it to address representational questions (Fernandino et al., 2016; Kay et al., 2008; Mitchell et al., 2008; Santoro et al., 2014; although some are more cautionary, e.g. Lee & Kuhl, 2016). While some authors have carefully restrained their discussion to the practical benefits of encoding models (Haxby et al., 2014; Davis & Poldrack, 2013), the interpretative benefits of encoding models over MVP classification are expressed strongly by many others. For example, in a recent article named “Resolving ambiguities in of MVPA Using Explicit Models of Representation”, Naselaris & Kay (2015) clearly state that:

> *With such models, it becomes possible to identify the specific representations encoded in patterns of brain activity and to map them across the brain. (p. 551)*

Similar strong claims were expressed by Ritchie et al (2017). They argued that by directly linking the structure of activation patterns to the structure of behavior, it becomes…

> …*implausible to suppose that the information is present but not correctly formatted, because the decoded format of the information in activation space is precisely what is being used to predict behavior in a psychologically plausible manner. (p. 23)*

A useful illustration of this inference in practice comes from a recent study by Fernandino et al. (2016). The authors wanted to understand how conceptual information is represented in a set of higher-order non-modality-specific brain regions in the General Semantic Network (Binder, Desai, Graves, & Conant, 2009). An encoding model based on subjective ratings for 5 sensory-motor features (“color”, “motion”, “sound”, “shape”, “action”) of training words was used to predict neural activation patterns related to novel individual words. The model successfully predicted above chance the brain activity patterns for concrete words in the semantic network regions (61% mean accuracy), but not in a set of control regions associated with visual word form processing. Based on these finding, Fernandino et al. (2016) suggested that *“the brain represents concepts as multimodal combinations of sensory and motor representations*” and that *“heteromodal areas involved in semantic processing encode information about the relative importance of different sensory-motor attributes of concepts, possibly by storing particular combinations of sensory and motor features”* (p. 9763).

Here lies the problem – this inference is not formally valid. We need to consider what the data would have looked like if the underlying neural representation was actually different (Mahon, 2015). In this example, the successful decoding of conceptual identity in the GSN based on an encoding model of sensory-motor features does not necessitate the representational format in the GSN to be sensory-motor in nature. The results might be obtained even if the GSN uses amodal representations, as long as there is a non-arbitrary mapping between representations in the GSN and sensory-motor features.

To illustrate, let us hypothetically assume that the GSN literally encodes word co-occurrence statistics. As co-occurrence statistics correlate with sensory-motor feature ratings, it would be possible to predict GSN activity patterns based on these features, even if they are not driving the activity patterns. Thus, while we can rule out the possibility that conceptual representations in heteromodal areas bear an arbitrary relation to sensory-motor features, as has been argued by some proponents of symbolic systems (Fodor & Pylyshyn, 1988), we cannot conclude that the GSN encodes multimodal sensory-motor information on the basis of Fernandino et al’s (2016) results. At most, we can say that the subjective sensory-motor relevance ratings that Fernandino et al. (2016) gathered for each concept capture some information represented in the GSN, but not whether this information is sensory-motor in nature.

This issue is not limited to the specific study discussed above. To put the claim more generally, we argue that information in one representational system might be decoded based on features from another, even if they use different representational schemes, *as long as there is at least a partially systematic mapping between them.* Specifically, while such encoding models should be able to predict the neural activation from the features of a stimulus if the brain uses a representational scheme based on those features (Naselaris et al., 2011), the reverse is not guaranteed1. A successful prediction can also occur when the stimulus feature space is *systematically related* to the features that underlie the neural representational scheme. However, that relationship need not be one of equivalence. There are at least three ways in which mappings between representational systems can be made and successful prediction can occur in two of those cases.

## V. Types of mappings

**Arbitrary mappings between representations.** First, items from two representational systems might be related in an entirely arbitrary way. For example, the meaning of words is mostly unrelated to their orthographic features2, and the geographic locations of countries are not predictive of their names, etc. More generally, consider two unordered sets of items, *A* = {*A*_1_, *A*_2_,…, *A*_*n*_} and *B* = {*B*_1_, *B*_2_,…, *B*_*n*_}. An arbitrary mapping between these two sets exists when the mapping from a specific item in set A to a corresponding item in set B is unrelated to the mappings between the remaining items in the two sets. In the context of encoding models and the brain, decoding of novel items from one set would be impossible based on a feature model from the other set, if these two sets are arbitrarily related.

**Sets that use the same representational scheme.** In contrast, a successful prediction can occur if the two sets use the same representational scheme. Consider the set of multi-digit numbers in the decimal system, *A* = {10,11,…,427,…}, and the set of 10 digits in the decimal system, *B* = {0,1,2,3,4,5,6,7,8,9,10}. These sets use the same representational scheme to represent quantities (the decimal system), and there is a systematic linear mapping from the features (the digits), to the multi-digit numbers, such that:

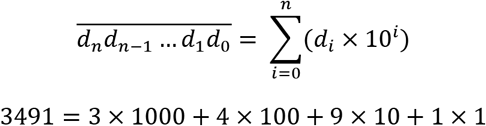

When we have such systematic mappings between systems that use the same representational scheme, knowing the mapping function allows us to decompose any item from set A as a combination of features from set B. An example of such a mapping would be Fernandino et al.’s (2016) suggestion that the General Semantic Network encodes multimodal combinations of sensory-motor features by integrating information from modality-specific sensory-motor areas. If this were true, then you could predict the neural pattern of novel items from their featural representations, which is what that study found as well.

**Sets that use different but systematically related representational schemes.** However, there is an alternative, which would also allow you to make a successful prediction from encoding models. Two sets can use different representational schemes, while at the same time maintaining a systematic mapping between themselves. That systematicity allows us to predict the mapping of any one pair of items from knowledge of the mapping function. Within the context of conceptual representations in the brain, higher-level heteromodal areas might use a representational code that is different from the one used by sensory-motor cortices, but there might be a systematic mapping between representations in each system3.

For a simplified example, consider the relation between the decimal and the binary systems for representing numeric values. A binary represented value can be transformed into a decimal number by applying the following formula:

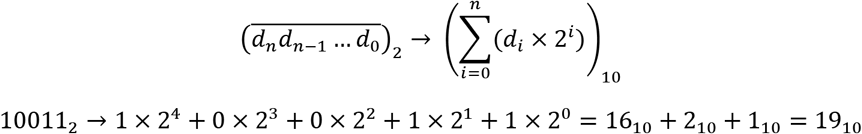

Clearly, there is a systematic but *non-linear* mapping between the decimal and the binary system, and yet, these two systems use different codes to represent numbers. If our argument is correct then it should be possible to predict the binary representation of a number based on a decimal feature encoding model. Below we present a simulation that achieves this by applying the encoding model approach often used in neuroimaging studies. Within the simulation, binary vectors are analogous to voxel activation patterns, and the encoding model is based on decimal representations (Table 1).

**Table 1.**
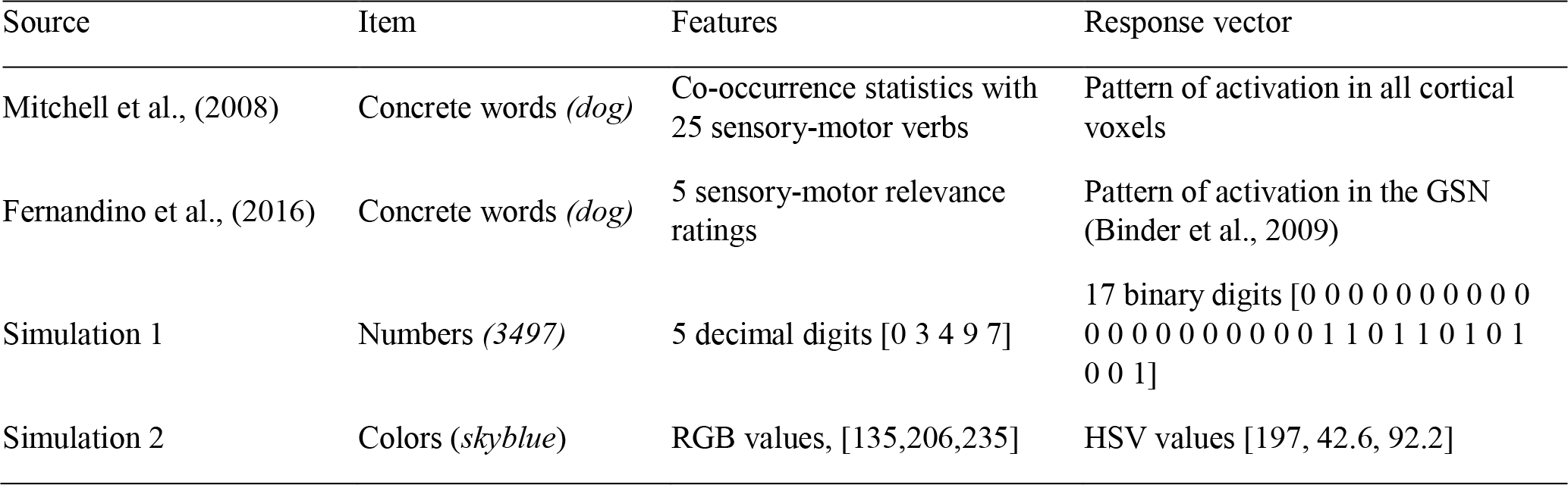
Examples of studies that use feature encoding models

## VI. Simulation 1: Decoding binary representations with a decimal feature encoding model

As detailed previously, encoding models predict stimulus identity from brain activation by modelling the relationship between the constituent features of the training stimuli and their corresponding BOLD activation in a group of voxels. Then they use that relationship to estimate the expected neural activation patterns for novel test items based on their feature representations. The predicted activation pattern for each stimulus is compared to the observed patterns for all test stimuli. For the following simulation, let us consider the numbers from 0 to 99 999 as our stimulus set. They can be decomposed into 5-dimensional feature vectors where each feature is a decimal digit (e.g., 3497 can be decomposed as [0 3 4 9 7]. These features can be considered analogous to the 5 sensory-motor relevance ratings of words used by Fernandino et al. (2016) or to the cooccurrence statistics with sensory-motor verbs used by Mitchell et al. (2008). Further, let us consider the binary representation numbers as 17-dimensional vectors (e.g. [0 0 0 0 0 0 0 0 0 0 0 0 0 0 0 0 0 0 0 0 1 1 0 1 1 0 1 0 1 0 0 1], to be analogous to the BOLD activation pattern in a set of 17 voxels in an ROI under investigation. The correspondence between these patterns and actual neuroimaging studies using this approach is demonstrated in Table 1.

We trained an encoding model to predict the binary activation pattern for a given number, based on its 5-dimensional decimal feature representation. The modelling followed 4 steps: 1) splitting the stimuli into a training (90%) and a test (10%) set, 2) fitting multiple linear regression models on the training set with the 17 binary features as response variables, and the 5 decimal features as predictors, 3) calculating predicted activation pattern (predicted maps, PMs) for each test item from its decimal features and the multivariate regression model, 4) comparing the PMs with the actual binary patterns for all test items (observed maps, OMs). In the comparison stage, we computed the Euclidean distance between each PM and the OMs for all test items, and we calculated the percentile rank of the similarity between the PM and the OM of each item. For example, if the PM for the number 29782 were most similar to the OM for that number, then the percentile rank for it would be 10 000/10 000 = 1. However, if it were more similar to the OMs of 1 000 other items, then its percentile rank would be 9 000/10 000 = 0.9.

The encoding model was successful in decoding the binary representation of untrained items based only on their decimal features. The prediction accuracy of the linear regression model was 0.7 (SD = 0.24) and a wilcoxon signed rank test showed that it was above chance (p < .0001). Since by definition binary and decimal number systems use different representational schemes, we cannot conclude that the representation of binary numbers encodes decimal features. By analogy, the successful decoding of patterns of neural activation based on a stimulus feature space, cannot be used to infer that the brain encodes information about these features or that its neural representational space is organized along the dimensions of that feature space.

## VII. Simulation 2: Decoding a color represented in one color space with an encoding model in another color space

Another way to illustrate how two systematically related, but distinct representational spaces can be decoded from one another, is to use color representational spaces. Any color can be uniquely defined as a point in a three-dimensional space. There exist numerous color spaces, and in some of them the dimensions reflect three distinct colors that can be mixed in different proportions to produce any other color (e.g., RGB, CMY), while in other color spaces the dimensions reflect attributes such as hue, saturation and lightness (e.g., HSL, HSV). RGB and HSV are both three-dimensional spaces, however, their dimensions are related non-linearly (see Figure 4). As a result, their representational geometry is different, and for example, one color can be equally distant from two others in RGB space, while it can be closer to one of them in HSV space. Despite this, the position of colors within those two spaces are systematically but non-linearly related, and there exist a number of algorithms to convert RGB representations to HSV and vice versa (for a primer on color spaces and color space conversions, see Ford & Roberts, 1998).

**Figure 4.**
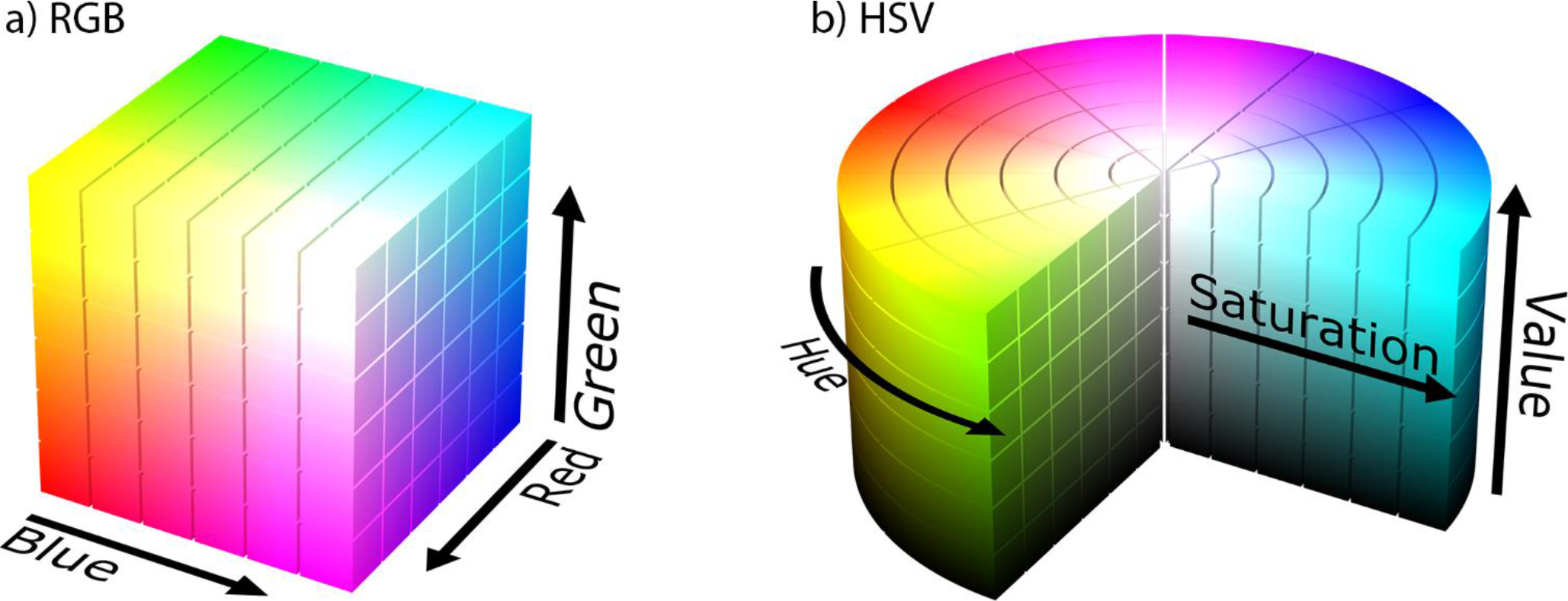
a) the RGB color space – black arrows show the three main color dimensions, whose values are subtracted to achieve the resulting color at each point in the color space; b) the HSV color space – black arrows show the value, saturation and hue dimensions that are combined to determine the color of each point in the color space. Images by Michael Horvath, available under Creative Commons Attribution-Share Alike 3.0 Unported license.

While we could build a complex encoding model that simulates neuronal tuning curves and voxel activations in each of these two spaces (similarly to Brouwer & Heeger, 2009), for illustrative purposes it would suffice to demonstrate that we can successfully predict the three valued HSV representation of novel colors, by training an RGB encoding model on a subset of randomly sampled colors. For the stimuli, we used 1000 colors by selecting 10 equally spaced values on each RGB dimensions. Thus, the stimulus features of each color were its three RGB values. The response vectors in this case were the corresponding three HSV values for each color. The decoding procedure was otherwise identical to the one in Simulation 1. We performed a 10-fold cross-validation. Similarly to Simulation 1, the encoding model based on RGB color representations was able to predict the HSV color representations of untrained colors quite well (mean rank accuracy 0.82, SD = 0.22, p < .0001).

The current simulation allows us to illustrate how such a high decoding accuracy in and of itself might not provide enough information to judge how well the encoding model reflects the neural representational space. For each item, we can directly compare the predicted color from the encoding model with the actual color of the stimulus. In Figure 5, we show the “observed” ground truth and the predicted color for each test item. Each panel shows hues for different combinations of saturation and value. Within each panel, the bottom half shows the actual color and the top half shows the predicted color based on the RGB encoding model. The model fares well in predicting the value and the saturation of the color. However, while overall it captures the hue progression, there are significant deviations. In summary, a high predictive accuracy is not by itself sufficient to conclude that an encoding model reflects the nature of the underlying neural representation.

**Figure 5.**
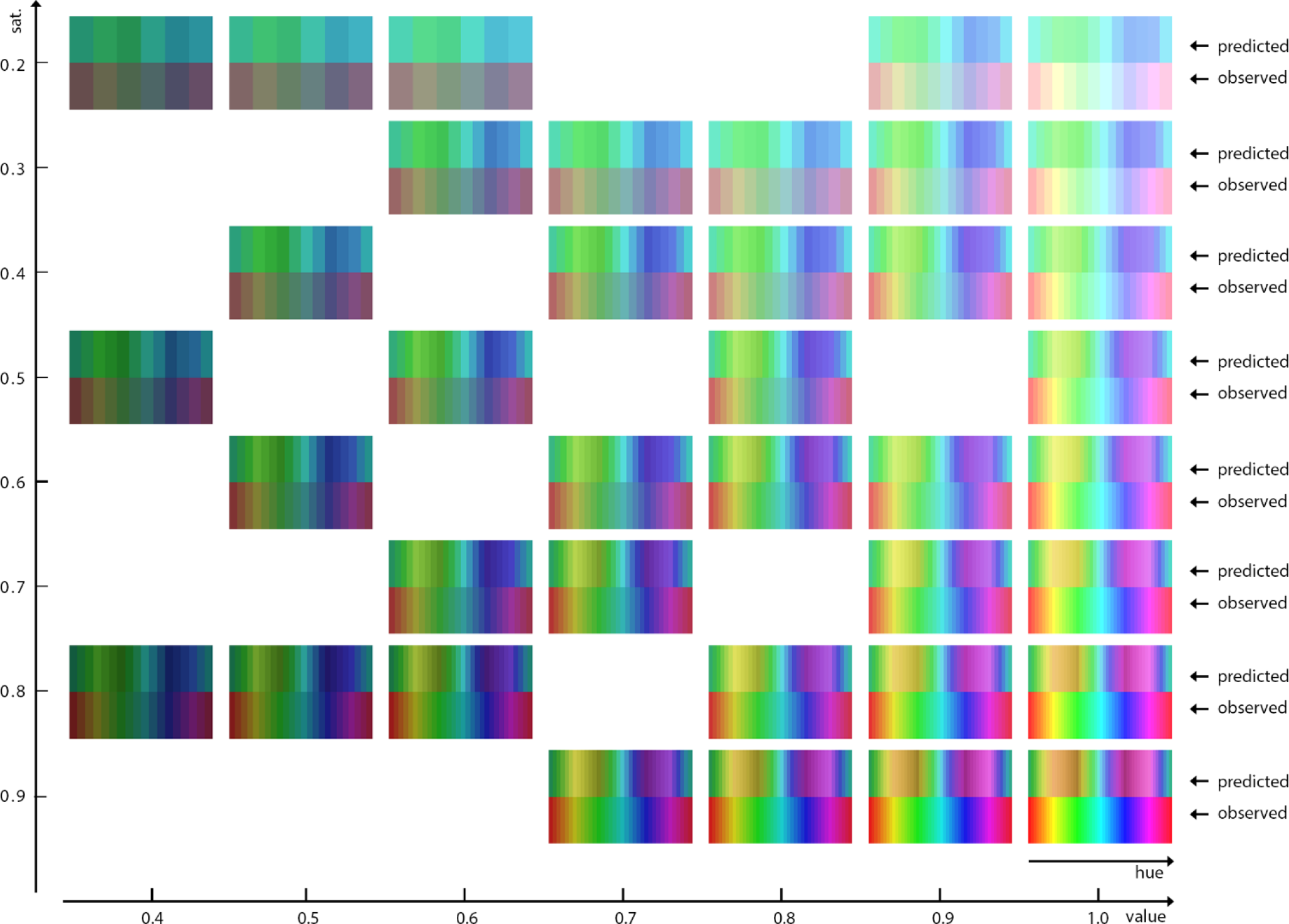
Results from fitting the RGB-based encoding model on the HSV color stimuli. Each mini-panel shows observed and predicted colors for a specific saturation and value combination. Within each panel, bottom row shows the sequence of HSV stimuli (i.e., ‘ground truth’ or ‘observed colors’), and the top row shows the predicted color from the RGB encoding model.

## VIII. Discussion

Stimulus-feature based encoding models (Haxby et al., 2014, Naselaris et al., 2011) are a powerful new tool for studying how the constituent features of stimuli relate to the neural activation patterns elicited by these stimuli. They represent a significant methodological advance over more traditional MVPA methods because they allow us to predict neural activation for novel items and because they can be used to decode the identity of such items from neural data alone. While this is an impressive feat and an incredibly useful tool, we have to be cautious in interpreting what such successes mean for our understanding of the representational system of the brain. Both theorists (e.g., Haxby et al., 2014; Naselaris & Kay, 2015; Naselaris et al., 2011; Norman et al., 2006; Tong & Pratte, 2012) and practitioners (e.g. Fernandino et al., 2016; Kay et al., 2008; Mitchell et al., 2008; Santoro et al., 2014) have suggested that we can infer that the brain uses a certain set of features to encode information, if we can successfully decode the activity of novel items from such features. However, as we have argued here, this inference is not formally valid. Successful decoding might be the result of a systematic relationship between the representational system of the brain and the stimulus feature set, even if those utilize different representational schemes.

### A. Representational equivalence – is it in the eye of the beholder?

How do we know whether two representational systems are truly different? It could be argued that in our first example, both binary and decimal number systems share many properties, and that they are merely different implementations of the same fundamental representation. For example, both systems use the position of a digit to encode its magnitude, and as a result, all arithmetic procedures that can be performed with decimal numbers can be applied to binary numbers as well. Despite these similarities, the transformation required to get the decimal from the binary representation of a number is non-linear. The point about linearity is important – it has been argued that two representational spaces could be considered equivalent only if there exists a linear transformation from one to the other (Naselaris et al., 2011). Linear transformations rotate multidimensional spaces, but do not change the nature of the representation, only the interpretation of the dimension. However, when the transformation required is non-linear, then the geometry of the underlying representational spaces is different (Kriegeskorte & Kievit, 2013).

We further propose that the key issue in determining whether two representational system are equivalent is whether you can establish a one-to-one mapping relation between features at different levels of representation in each system. For example, if you substitute each decimal digit with a unique letter, the resulting system would appear to be very different from the decimal system only on the surface – the relation between multi-digit numbers and their features would be the same in both cases1. In contrast, decimal and binary features have a qualitatively different relation to the numbers they represent. Despite this, binary representations can be decoded based on decimal features, illustrating the inferential problem of encoding models we address here.

Simulation 2 allowed us to illustrate this point more clearly by directly visualizing the differences in the geometries of RGB and HSV color spaces. While both color models use 3 dimensions to represent colors, these dimensions have distinct interpretations and can be related by a systematic non-linear transformation. The RGB color model is an additive mixture space, in which any color is produced by mixing the three primary colors in different proportions. In contrast, the HSV model represents colors by specifying their hue (e.g., red), their saturation (i.e., the intensity of the pigment), and their value (i.e., the darkness of the pigment). These three-dimensional representational spaces have different geometries, such that the relative distances between colors are not identical. Yet, training an encoding model to predict the HSV representation of novel colors based on the relationship between the RGB and HSV features of other colors, was highly accurate on average.

### B. Where to go from here?

#### 1. Model comparison

An important question that naturally arises from the caveats we discussed is how one can maximize confidence in the outcome of a forward encoding model approach, or conversely, guard oneself against unjustified inferences. As others have noted, it is crucial to compare the performance of several possible encoding models (Haxby et al, 2014; Naselaris et al., 2014). However, to that aim, it is not sufficient to use a "baseline model" that is unrelated to the domain of interest (i.e., comparing a semantic feature model to a low-level visual word form model as exemplified by Fernandino et al., 2016). Instead, one or several alternative representational models should be tested that are derived from competing theories (i.e., semantic model A vs. semantic model B). To illustrate, an elegant comparison of a sensory-based vs. non-sensory-based semantic model was achieved by Anderson et al. (2015). These authors contrasted a visual model with a word co-occurrence model to investigate which brain regions represent modality-specific visual features, and which do not (using differential correlation in RSA rather than an encoding model). The relative superiority of a particular model at predicting activation patterns in a brain region makes it more likely that the brain is using the representational scheme of the better performing model rather than the alternative. However, it is important to keep in mind that such comparisons only provide evidence for the relative likelihood of each model, but, due to the limitations discussed above, still do not allow us to infer that the winning model is the “true” model (Palminteri, Wyart, & Koechlin, 2017).

#### 2. Absolute model performance

For that reason, besides the assessment of relative model performance based on model comparison, a second crucial step is to evaluate absolute prediction performance. In particular, the observed decoding accuracy can be compared to the “noise ceiling”, or to the “upper limit of prediction accuracy” (Naselaris et al., 2011), reflecting the maximal performance that can be feasibly achieved given the noise present in the signal. The gap between the two can be thought of as the variance that is not explained by the current model, which should motivate and guide the search for an improved or alternative version of the model. Until such maximal performance is obtained, we should be careful in making strong representational inferences about the brain from the currently available analytic methods.

#### 3. Beyond the decoding accuracy: visualizing and comparing the organization of predicted and observed neural activation patterns

It is important to note that even high decoding or predictive accuracy on its own is insufficient to establish representational equivalence. Despite the fact that in Simulation 2 our RGB encoding model predicted the HSV representation with an 82% rank accuracy, a more careful inspection of the actual and the predicted HSV colors revealed that there are significant deviations in the model predictions. This is because the regression model is attempting to establish a linear rotation between the non-linearly related RGB and HSV spaces.

This result should make us weary of making conclusions of representational equivalence based on a single accuracy value, as is the practice in some published studies (e.g. Fernandino et al., 2016). Rather, it is important to examine the actual correspondence of the representations with additional RSA methods or dimensionality reduction and visualization techniques (e.g., Brouwer & Heeger, 2009; Foster et al., 2016). For example, rather than depending on decoding accuracy alone, Brouwer & Heeger (2009) used principle component analysis (PCA) on the predicted voxel activation patterns and showed that the two main components present in the signal corresponded well to the actual organization of the colors in the color space they were using. Similarly, Foster et al. (2016) showed that activity in the hypothesized angular channels does indeed follow a graded pattern that peaks at their preferred orientation. In summary, we believe it is crucial to carefully examine the model predictions. If the encoding model reflects the actual representational space, there should be no systematicity in the errors it makes, and the *relations* between the predicted activation patterns should resemble the relations and the organization of the observed activation patterns.

#### 4. Attentional modulation

Ultimately, many of these inferential caveats exist because fMRI data is correlational. Comparing alternative models and evaluating absolute prediction performance might eventually converge on the true underlying feature model, but this is not guaranteed. We propose that an even better way to test representational hypotheses might be to experimentally manipulate the hypothesized representational dimensions. For example, one could prime participants to weight or attend to some features of the stimuli more than others. This type of attentional modulation decreases noise in the correlated activity between individual neurons, which enhances population coding (Downer, Niwa, & Sutter, 2015), and it also increases selectivity of individual neurons for task-relevant stimulus features (Sigala & Logothetis, 2002). Thus, if orienting attention to some features of the stimuli improves the ability of an encoding model based on those features to predict neural activity, this would constitute much stronger evidence for the viability of the model.

Relatively little work exists to evaluate this proposal with fMRI, although two recent studies present promising results with an attentional modulation method that could eventually be extended to encoding model analyses as well. Çukur, Nishimoto, Huth, & Gallant (2013) showed that when participants were searching for people or for vehicles in movie clips, individual voxels became more attuned to the attended category. This occurred regardless of whether the target category was actually present in a scene. Importantly, this tuning shift was progressively stronger in higher-order visual areas compared to early retinotopic visual areas. Thus, attentional tuning likely reflects the fact that some aspect of the attended information is being represented in a specific ROI.

Even more relevant is a recent study by Nastase et al., (2017), who extended this approach to representational similarity analysis. Participants saw a sequence of short clips with animals performing certain behaviors, and they had to respond to each clip in one of two ways. In one condition, the task was to say whether the animal in the current clip is from the *same taxonomic category* as the animal in the previous clip. In the other condition, participants had to respond whether the animal was performing the *same behavior* as the animal in the previous clip. The authors derived two representational dissimilarity matrices for the stimuli – one based on the taxonomic dissimilarity, and one based on the dissimilarity in the behavior that the animals performed. They calculated the correlation of these two representational dissimilarity matrix (RDM) with an RDM derived from the neural signal. When participants attended to the taxonomic category of the animals, the correlation between the neural and the taxonomic RDM increased. In contrast, attending to the animal’s behavior increased the correlation of the neural RDM with the behavioral RDM.

Perhaps the most relevant finding of the study is that even though there was a significant correlation between the model RDMs and the neural RDMs in several regions, attentional modulation *did not increase* the correlation in all of them. For example, while the behavioral model correlated with neural activity in both the early visual cortex, and in post-central parietal regions involved in motion perception, attending to the behavior of the animals increased the correlation only in the latter regions. Thus, on the basis of the overall correlation one might have concluded that behavior is represented even in early visual cortex; however, the lack of the attentional modulation in those areas indicates the overall correlation might have been driven by some visual information that is simply confounded with the animal's behavior. Similarly, taxonomic information correlated with the early visual cortex and the ventrotemporal cortex, but attending to taxonomy increased the correlation only in the latter. Overall, these results provide some indication that modulating attention to specific features of the stimuli can be beneficial in determining whether successful decoding reflects the nature of the neural representation, or whether it is picking up systematically related information instead. We expect that this technique can be effectively extended to encoding models as well.

### C. Final remarks

Many of the points we have raised here are not specific to encoding models, but can also be leveled against any other method currently in use for understanding neural representations. The reason why we focused here on encoding models is that they are becoming increasingly popular in recent years, and we believe it is important to highlight their limitations. The speed at which decoding methods have developed since Haxby et al.’s (2001) seminal study on multivariate analyses represents the importance of these methods to the neuroscientific community. Encoding models have many advantages over more traditional MVPA techniques, such as the ability to decode items that the model has not been trained on. There is no doubt that this represents a great technological advance. However, exactly because encoding models are so powerful, it is important to understand what inferences we can and cannot draw from them. There is little doubt that successful decoding in a certain brain area tells us a lot about *where* in the brain the decoded information might be represented. The question of how those representations are organized, albeit, is more difficult and cannot be answered only on the basis of a single significant decoding value. Our hope with this commentary is not to downplay the important role of these methods, but to further the discussion about what we can and cannot learn by using them.

## Footnotes

1 this problem is similar, but not identical, to the problem of reverse inference (Poldrack, 2006)

2 Whereas a minor degree of systematicity does seem to exist in this domain (e.g., Monaghan et al., 2014), word meanings cannot be systematically predicted based on their orthography and vice versa.

3 What makes representational codes different is a surprisingly difficult question to answer. Due to space limitations we will briefly cover this issue in the general discussion, but a more in-depth treatment is needed

4 in fact, because of that linear one-to-one relationship, replicating our simulation with these two examples leads to perfect decoding accuracy; compare that to the 0.7 decoding accuracy for the decimal-to-binary model

